# Contributions of superior colliculus and primary visual cortex to visual spatial detection in freely moving mice

**DOI:** 10.1101/2025.09.11.675538

**Authors:** Jisoo Kim, Riccardo Beltramo, Jasper Poort

## Abstract

Visual spatial detection is a crucial first step in vision to guide decision-making and action. However, its neural basis in freely behaving animals remains unclear due to challenges in controlling visual input and monitoring eye and head position. Studies in head-fixed mice have shown that both the superior colliculus (SC) and primary visual cortex (V1), two key visual processing hubs in mammals, contribute to visual spatial detection. Yet, their relative roles in freely moving animals are poorly understood. Here, we developed a novel approach to study the neural mechanisms of visual spatial detection in unrestrained mice. We combined closed-loop presentation of visual stimuli with neural recordings, optogenetic manipulation, and simultaneous monitoring of eye and head position. Our results show that SC cells are more predictive of reaction times than V1 cells. Furthermore, SC neurons exhibit more sustained activity than V1 neurons during visual spatial detection. Optogenetic SC and V1 silencing causes pervasive and remarkably localized perturbations of visual detection performance. SC silencing has a stronger impact on visual detection than V1 silencing. These results highlight the distinct activity patterns in two principal early visual processing centers, and establish their relative causal contributions to visual spatial detection.

## Introduction

Detecting visual stimuli and responding appropriately is essential for survival in natural environments. The primary visual cortex (V1) and the superior colliculus (SC) are two key hubs in visual processing that belong to two major distinct visual pathways and contribute to visual spatial detection (Basso et al., 2021; Beltramo, 2020; Beltramo and Scanziani, 2019; Cone et al., 2024; Glickfeld et al., 2013; Hafed et al., 2023; Krauzlis et al., 2013; Lee et al., 2020; Wang et al., 2020, 2010; Wheatcroft et al., 2022; White et al., 2017). Most of what we know about their contributions comes from studies in head-fixed animals, where visual input can be precisely controlled and behavior closely monitored. However, in the natural world, animals rarely experience vision while immobile. Instead, they continually move their eyes, head, and body to sample the environment, detect threats, locate food, and guide action. Thus, understanding visual detection under naturalistic conditions is essential, because eye and head movements shape visual input, modulate neural responses, and engage circuits that may operate differently compared to restrained settings. Growing interest in naturalistic and ethological behavior (Dennis et al., 2021; Franke et al., 2024; Gantar et al., 2025; Hoy and Farrow, 2025; Klioutchnikov et al., 2023; Lanzarini et al., 2025; Liu et al., 2016; Qiu et al., 2021; Shapcott et al., 2025; Skyberg and Niell, 2024; Wallace and Kerr, 2019) has led to the development of methods for monitoring eye and head movements in freely moving animals (Meyer et al., 2020, 2018; Michaiel et al., 2020; Parker et al., 2022; Payne and Raymond, 2017; Singh et al., 2025). Studying visually guided behavior in freely moving mice offers the opportunity to capture more active and engaged visual processing and to examine the full range of eye and head movements (Meyer et al., 2018, 2020; Parker et al., 2022, 2023; Payne and Raymond, 2017; Sawinski et al., 2009; Sharp et al., 2025.; Singh et al., 2025). Despite these advances, how SC and V1 contribute to visual spatial detection under freely moving conditions remains largely unknown.

Here, we present a method to study the neural mechanisms of visual spatial detection in freely behaving mice by combining closed-loop visual stimulus presentation with neural recordings, optogenetic manipulation and eye and head position tracking. We first confirmed that we were able to observe spatially selective visual responses in both SC and V1, validating our approach for studying visual processing in freely moving mice. We then examined how SC and V1 neural activity correlated with task performance. Early SC activity was more strongly correlated with reaction time than early V1 activity. Beyond the early period, SC showed sustained neural firing, whereas V1 activity was more transient. Finally, to test causality, we used optogenetic manipulations and found that SC inhibition impaired task performance more than V1 inhibition. Together, these findings establish a framework for investigating visual spatial detection in freely moving mice and show that the SC contributes more than V1 to visual spatial detection.

## Results

### Closed-loop visual detection task combined with neural recordings in V1 and SC

There are two major obstacles to understanding how freely moving animals process visual information— control of visual input and measurement and perturbation of neural activity during dynamic behavior. We therefore developed an integrated system combining closed-loop stimulus presentation and a head-mounted system for behavioral and neural measurement and optogenetics (Figure 1A). The closed-loop system controlled visual input by presenting visual stimuli only when the mouse faced the screen (Figure 1B and S1C-1D). Each trial started when the mouse entered the region of interest (ROI) and its head angle relative to the touchscreen was below 45 degrees. The head-mounted system, adapted from previous work (Meyer et al., 2020), enabled simultaneous behavioral and neural recordings with optogenetic manipulations in SC and V1 (Figure 1A and S1A).

**Figure 1:**
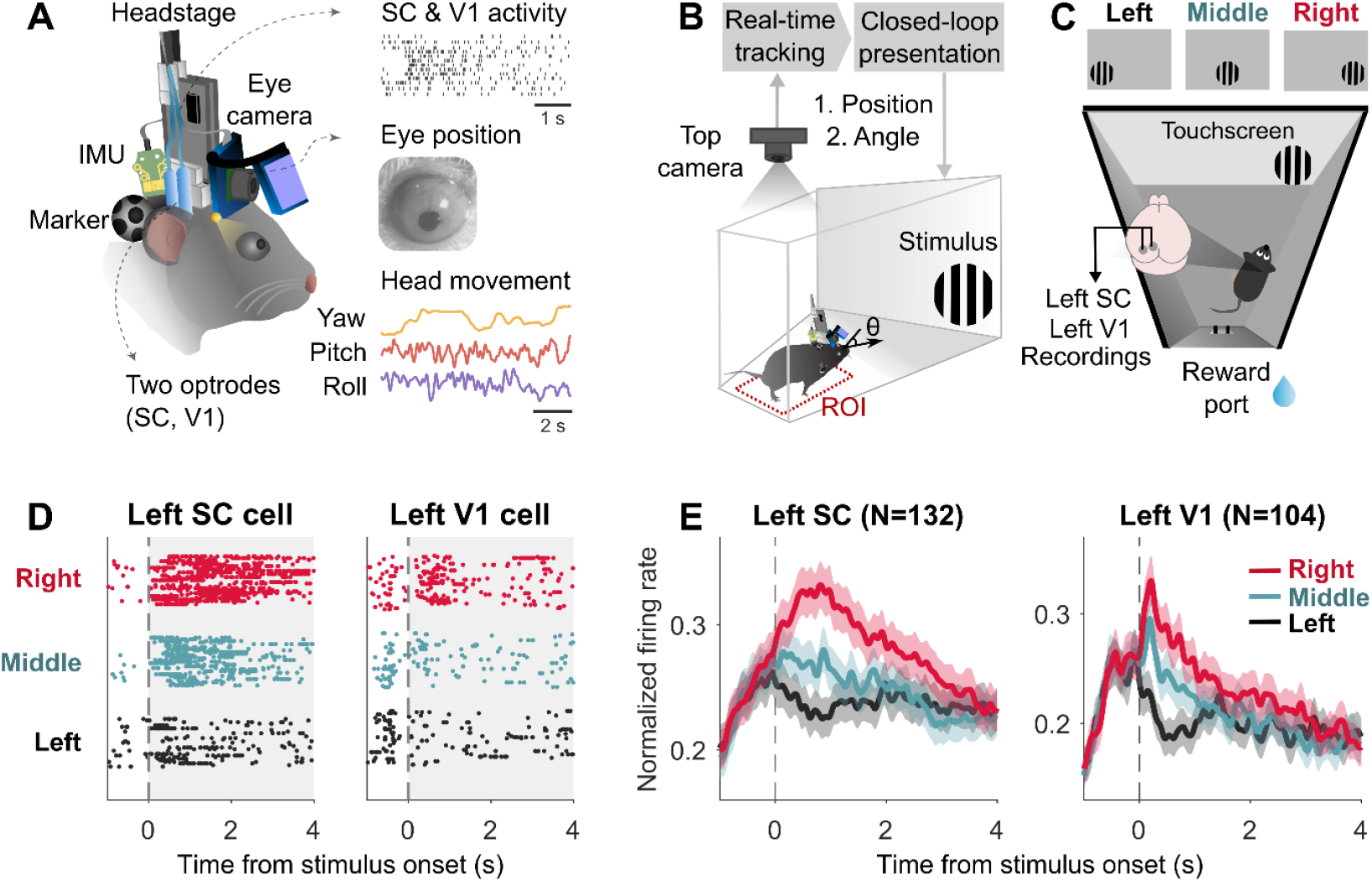
Visual responses in SC and V1 during a closed-loop detection task in freely moving mice. (A) Integrated head-mounted system with eye camera, inertial motion sensor (IMU) and markers for behavioral measurement and optrodes for neural recording and optogenetic inhibition. Two optrodes (tetrodes with optic fiber) were implanted in the left V1 and SC (16 tetrodes, 8 in each area). Head-mounted miniature eye camera monitored right eye position (see Figure S1A and S1B). IMU measured head movement (pitch, roll, yaw). Styrofoam markers were used for tracking the head position. (B) Schematic of closed-loop visual stimulation in freely behaving mice. Top camera measured the location of the mouse in real time. The visual stimulus was only presented when the mouse was in the region of interest (ROI) and faced the touchscreen (see Figure S1C and S1D). (C) Visual detection task in freely behaving mice (see Figure S1E). A drifting grating stimulus was displayed on the touchscreen when the mouse was in the ROI and faced the touchscreen. This stimulus appeared randomly in one of three locations: left, middle, or right. Mice received strawberry milk rewards from a lick spout when touching the target stimulus. (D) Raster plots of example SC (left) and V1 neurons (right). Each line represents a spike. Visual stimuli were presented at t = 0 s at three spatial locations (black: left stimulus, green: middle stimulus, red: right stimulus, see Figure S1F). (E) Normalized average firing rates of SC neurons (n = 132 neurons from 6 mice) and V1 neurons (n = 104 neurons from 7 mice). Shaded area represents SEM. Normalized firing rates were calculated by dividing each cell’s firing rate by its maximum value. Note the increase in firing activity before stimulus onset, as the animal orients its head towards the screen.

Visual responses in SC and V1 during visual detection tasks are well characterized in head-fixed conditions (see for example Cone et al., 2024; Glickfeld et al., 2013; Ito and Feldheim, 2018; Wang et al., 2020). Using our integrated system, we confirmed spatially localized visual responses in SC and V1 in freely moving mice. We simultaneously recorded single-unit activity from superficial layers of SC and V1 neurons during a spatial detection task. Visual stimuli (horizontally drifting gratings) were presented at three different spatial locations (Figure 1C). As expected from left hemisphere recordings, neurons in SC and V1 exhibited significantly stronger normalized firing rates for right (contralateral) stimuli compared to left (ipsilateral) stimuli (Figure 1D-1E and S1F; left vs right stimulus, SC: p = 1.25 × 10^−22^, V1: p = 1.09 × 10^−17^, Wilcoxon signed-rank, Bonferroni-corrected).

### Early SC Activity Predicts Reaction Time Better Than V1 During Visual Detection in Freely Moving Mice

Next, we investigated the relation between task performance and SC and V1 activity. Specifically, we determined whether early SC and V1 activity differentiated between fast and slow behavioral responses during visually-guided spatial detection in freely moving mice (Figure 2A). Previous work in head-fixed mice has shown that SC and V1 activity correlates with behavioral responses (Cone et al., 2024; Resulaj et al., 2018; Wang et al., 2020). Here, we defined reaction time (RT) as the time from stimulus onset until the mouse approached and touched the correct stimulus. Based on the reaction time, trials were categorized as ‘fast’ or ‘slow’ trials (fast: RT < 2.5 s; slow: 2.5 s < RT <10 s). SC neurons showed significantly higher firing rates during the early time window (0-1 s) in fast trials compared to slow trials. V1 neurons also showed a significant difference, but with a smaller effect size (Figure 2B and 2C, Miss vs Fast Hit, SC: p = 5.08 × 10^−10^, V1: = 1.45 × 10^− 3^, Wilcoxon signed-rank). Comparing the difference between fast and slow trials in SC and V1 showed a significantly larger difference in SC (Figure S2C, Wilcoxon rank-sum test, p = 0.0019).

**Figure 2:**
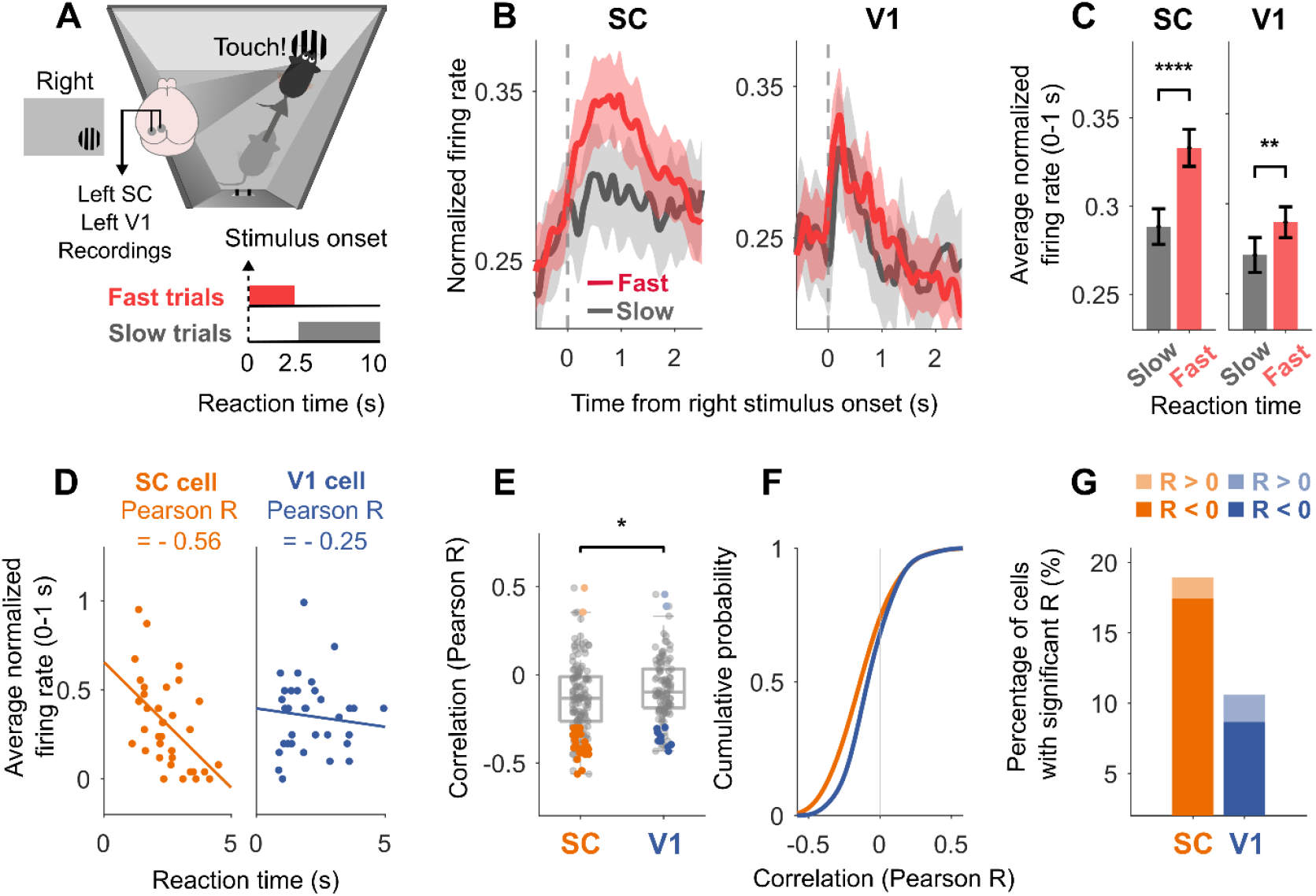
Early SC neural activity predicts reaction time during visual detection better than V1. (A) Schematic diagram of behavioral categories. Reaction time (RT) was calculated from stimulus onset to the moment the mouse touched the drifting grating. Trials were classified into two behavioral categories based on RT: fast trials (RT < 2.5 s) and slow trials (2.5 s < RT < 10 s). (B) SC and V1 neural activity in fast (red) and slow trials (grey). Shaded area represents SEM. (C) Average SC and V1 activity during the early period (0–1 s). Error bars, SEM. (D) Correlation between reaction time and early stimulus response (0-1 s) in SC (left) and V1 neuron (right). Each point represents an individual trial. (E) Correlation coefficients of SC and V1 all neurons (neurons with significant negative correlation; SC: orange, V1: blue; neurons with significant positive correlation; SC: light orange, V1: light blue). (F) Cumulative distribution of correlation coefficients of SC (orange) and V1 neurons (blue). (G) Percentage of cells with significant negative (SC: orange, V1: blue) or positive (SC: light orange, V1: light blue) correlation in SC and V1 (SC: 19%; V1: 10%).

To further investigate the relationship between early neural activity and reaction time, we analyzed trial-by-trial correlations in individual neurons. We calculated the correlation between reaction time and early activity (normalized firing rate from 0-1 s) for each SC and V1 neuron (Figure 2D). SC neurons showed significantly more negative correlations between reaction time and early activity than V1 neurons (Figure 2E-2F, Wilcoxon rank-sum test, p = 0.018). This pattern remained consistent when restricting head angle or excluding trials with reaction times under 1 second (Figure S2D-2P). We also calculated the percentage of neurons with significant correlation between their activity and reaction time. The majority of these cells exhibited negative correlations, meaning that higher early firing rates were associated with shorter reaction times. Notably, SC contained a higher proportion of significant cells than V1 (Figure 2G, SC: 19%, V1: 10%). These results show that SC activity during the early phase of stimulus processing is more predictive of reaction time than V1.

### SC Exhibits Sustained Firing Activity Throughout Visual Detection, While V1 Displays Transient Responses

Beyond the early phase of activity, how do SC and V1 responses differ during the visual detection task? Strikingly, we observed that a subset of SC neurons fired continuously from stimulus onset until the mouse touched the correct stimulus (Figure 3A and Figure S3D). In contrast, V1 neurons typically exhibited transient responses around stimulus onset. To analyze these patterns across the population, we aligned SC and V1 activity to the time of first touch to the correct stimulus (Figure 3B). SC neurons showed elevated and spatially localized firing during the pre-touch period. The difference in pre-touch activity between left and right stimuli was significantly greater in SC than in V1 (Figure S3C, Wilcoxon rank-sum test, p= 0.00099, SC: 110%, V1: 54%).

**Figure 3:**
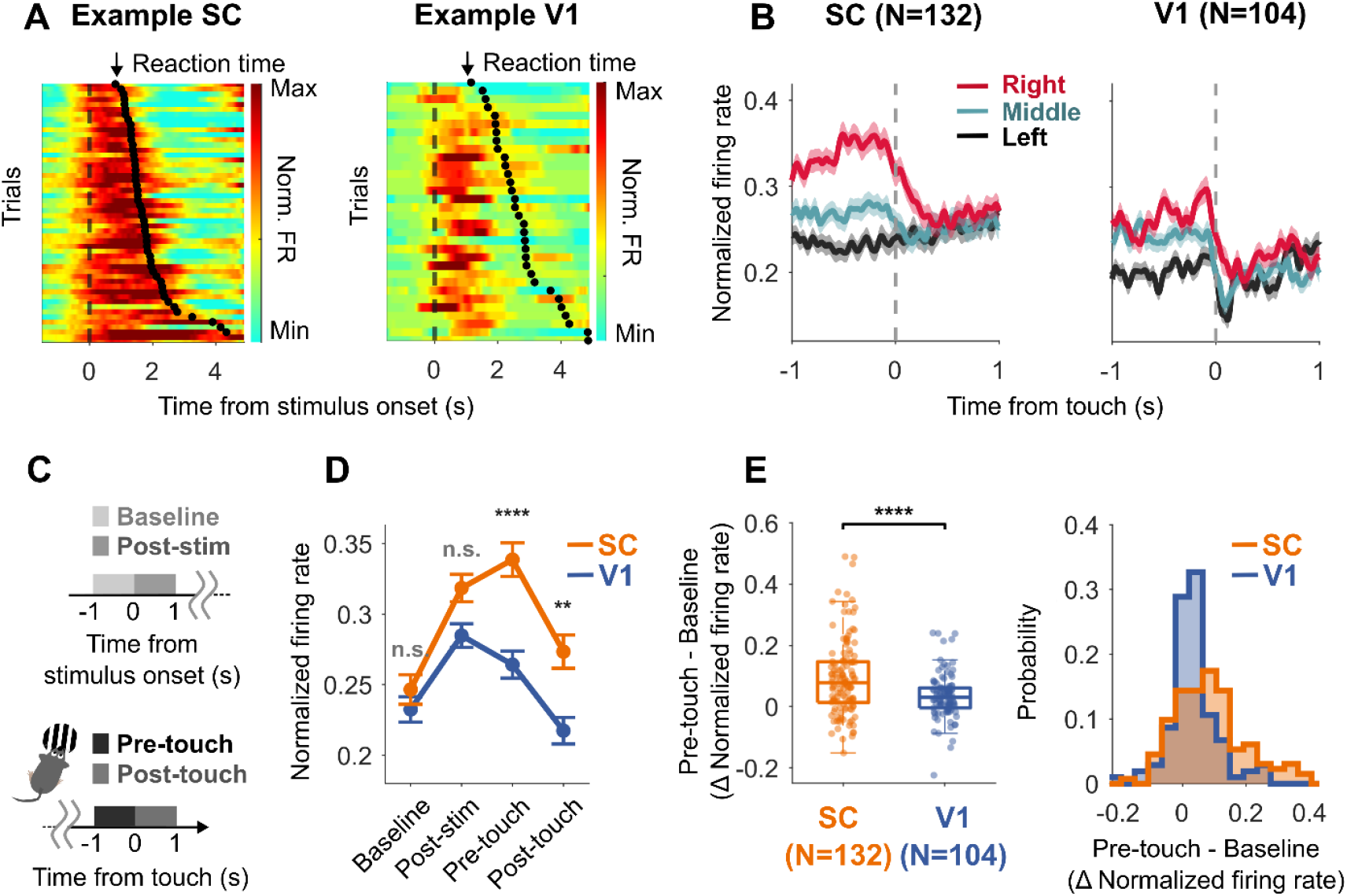
Sustained SC firing Activity Compared to Transient V1 Responses. (A) Example of visual responses of individual SC and V1 neurons. The mean baseline firing rate (–1 to 0 s) was subtracted from normalized firing rates for each cell. Black dots mark the reaction time for each trial, defined as the moment the mouse touched the screen, and trials are sorted from shortest (top) to longest (bottom) reaction time. The SC neuron exhibited sustained activity, firing from stimulus onset until the mouse touched the stimulus, whereas the V1 neuron showed a transient response. (B) SC and V1 activity aligned to touch onset, defined as the first correct touch. Black, green, and red lines indicate left, middle, and right stimuli, respectively. Shaded area represents SEM. (C) A schematic diagram of four phases throughout the detection task, Baseline (−1 to 0 s before stimulus onset), Post-stimulus (0 to 1 s after stimulus onset), Pre-touch (−1 to 0 s before touch onset), and Post-touch (0 to 1 s after touch onset). (D) Temporal dynamics of SC and V1 across the task. SC showed higher pre-touch activity than V1. Error bars, SEM. (E) The difference between Pre-touch and Baseline activity in SC and V1. (Left) Scatter plot showing the difference between Pre-touch and Baseline activity in SC and V1. (Right) Probability distribution of the difference between Pre-touch and Baseline activity in SC and V1, revealed that SC showed significantly higher pre-touch activity compared to V1 (SC: orange, V1: blue).

We next characterized how SC and V1 firing activity evolved throughout the detection task. To quantify temporal dynamics across the task, we divided the detection task into four phases (Figure 3C and Figure S3A), Baseline (−1 to 0 s before stimulus onset), Post-stimulus (0 to 1 s after stimulus onset), Pre-touch (−1 to 0 s before touch onset), and Post-touch (0 to 1 s after touch onset). First, during the post-stimulus period, both SC and V1 exhibited increased firing rates relative to baseline, reflecting the expected visual responses (Figure 3D). However, during the pre-touch phase, SC firing rates were significantly higher than those in V1 (Figure 3D, Wilcoxon rank-sum test, p = 1.5 × 10^−5^, Bonferroni-corrected). The difference between Pre-touch and Baseline was significantly greater in SC than V1 (Figure 3E, Wilcoxon rank-sum test, p = 0.0001). Thus, SC exhibited more sustained firing activity than V1 throughout the visual spatial detection task.

### SC Inhibition Impairs Visual Spatial Detection More Than V1 Inhibition in Freely Moving Mice

Since we found that SC showed stronger correlations with reaction time and displayed more sustained firing activity than V1, we hypothesized that SC contributed more to visual detection performance than V1. We therefore selectively inhibited SC or V1 to causally test the effects on the behavior task. We optogenetically activated local GABAergic neurons (Beltramo and Scanziani, 2019; Senzai et al., 2019; Wang et al., 2020) to suppress activity in the left SC or V1. The inhibition reduced neural activity in both areas to a similar extent (Figure 4B-4C, and S4A-C; SC: −40%, V1: −37%, p = 0.48). To determine whether V1 and SC inhibition caused motor impairment, we compared eye and head movements across conditions (control, V1 inhibition, and SC inhibition). Head orientation (roll, pitch, and yaw) and eye positions (horizontal and vertical) were comparable during the early phase (0-0.5 s) under V1 inhibition and SC inhibition compared to control condition (Fig 4D-4G, all n.s., see S4D-I).

**Figure 4:**
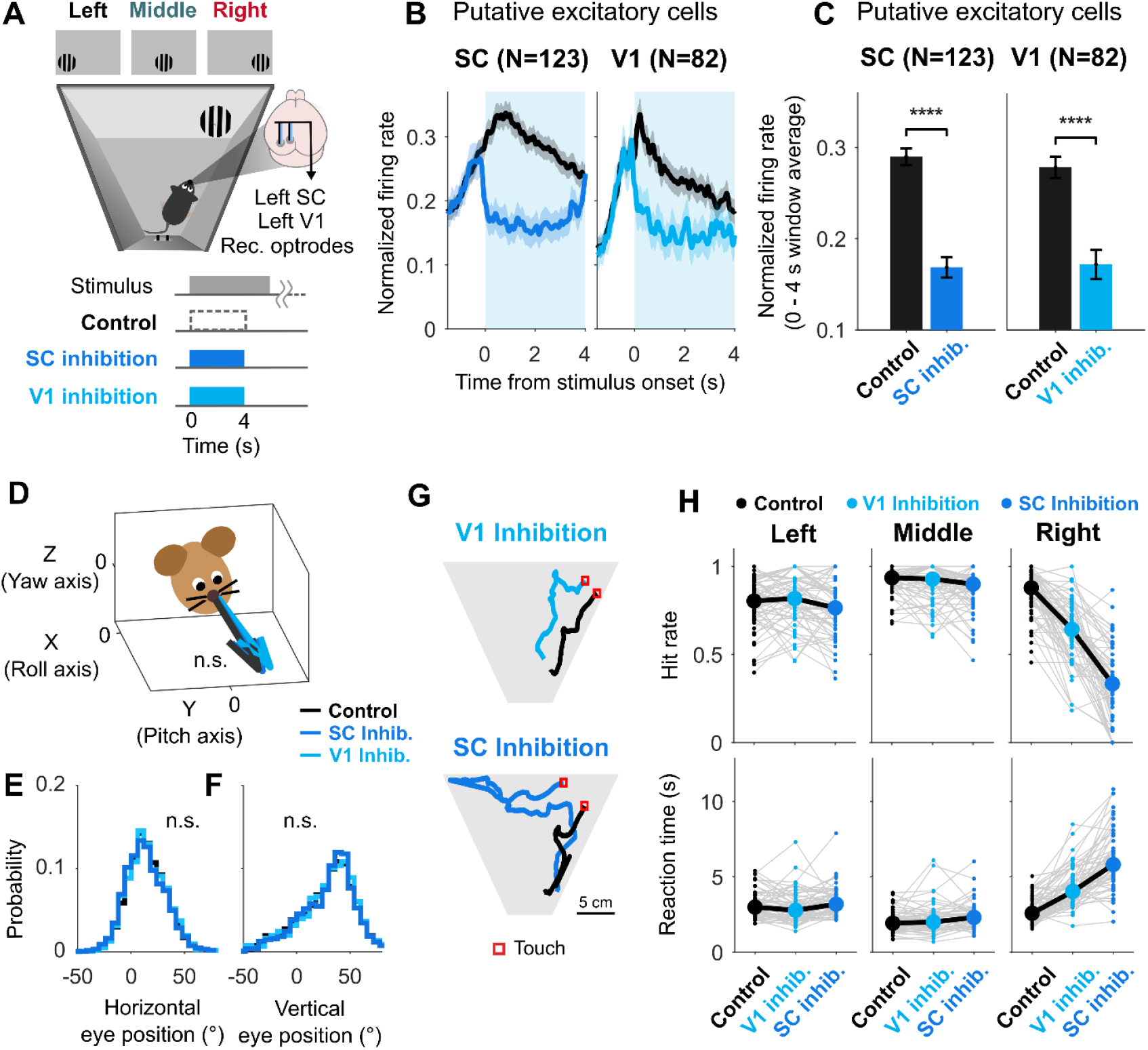
Impact of Optogenetic Inhibition of V1 and SC on Visual Spatial Detection in Freely Moving Mice. (A) Visual detection task in freely behaving mice with optogenetic silencing of V1 or SC. Visual stimuli were presented at three locations. In the control condition, no optogenetic inhibition was applied. During V1 or SC inhibition, activity was silenced for 4 seconds after visual stimulus onset. (B) SC and V1 activity in response to right stimulus during local optogenetic inhibition. Black line represents control condition and blue line represents local optogenetic inhibition (left: V1 activity during V1 inhibition, right: SC activity during SC inhibition). Shaded area represents SEM. Optogenetic silencing lasted for 4 seconds from stimulus onset. (C) Average normalized firing rate of SC and V1 (0-4 s). Error bars, SEM. (D) 3D head orientation vectors across conditions (control: gray, SC inhibition: blue, V1 inhibition: light blue) during the early period (0-0.5 s). Arrows indicate the mean head-direction vector for each condition, computed by rotating a unit vector according to the average yaw, pitch and roll angles within this period. Vectors are shown in a head-centered coordinate system, with the X (roll axis), Y (pitch axis), and Z (yaw axis) (see Figure S4E-4G.) (E-F) Horizontal and vertical eye position across conditions (control: gray, SC inhibition: blue, V1 inhibition: light blue, see Figure S4H-4I). (G) Trajectory of mouse during example trials in visual detection task (black line: control, blue line: inhibition condition). (H) Impact of optogenetic inhibition on behavioral performance, including hit rate (top) and reaction time (bottom). See main text for quantification of percentage change and statistical comparison.

We quantified behavioral performance using the hit rate (defined as the percentage of correct responses within 4 seconds). Inhibiting either V1 or SC led to a significant reduction in hit rate compared to the control condition, indicating that both areas are involved in visual detection (Figure 4G-4H). Inhibition was applied only to the left hemisphere, and performance was primarily impaired for right (contralateral) stimuli, as expected. Specifically, the hit rate for the right stimulus dropped by 26.2% under V1 inhibition (p = 1.5×10^-7^) and by −59.3% under SC inhibition (p = 8.7×10^-9^), while changes in hit rate for left stimuli were minimal (SC: −6.5%, p = 0.027; V1: 1.8%, p = 1, Bonferroni corrected, Wilcoxon signed-rank). While both SC and V1 inhibition impaired detection performance, the reduction was significantly greater with SC inhibition compared to V1 inhibition (p = 2.4×10^-8^, Wilcoxon signed-rank, Bonferroni corrected). Similarly, inhibition of SC and V1 increased reaction time (V1: p = 1.4×10^-8^, SC: p = 7.5×10^-10^, Bonferroni corrected, Wilcoxon signed-rank), and the effect was stronger for SC (p = 6.4×10^-8^, Bonferroni corrected, Wilcoxon signed-rank).

## Discussion

To dissect the neural underpinnings of visual spatial detection in freely behaving mice, we developed an integrated approach that combines optogenetic manipulations and closed-loop control of visual stimuli with simultaneous measurements of neural and behavioral activity (Figure 1). This method allowed us to probe visual processing under naturalistic conditions while retaining experimental control. Our results demonstrate that SC contributes more strongly than V1 to visual spatial detection in unrestrained mice. We found that SC firing activity during the early response window after stimulus presentation was more predictive of reaction times than V1 activity (Figure 2). Beyond the initial response window, SC showed markedly sustained neural firing, whereas V1 activity was more transient. Notably, a subset of SC neurons exhibited prolonged activity tightly aligned with the moment the mouse hit the visual target (Figure 3). Finally, causal manipulations with optogenetic silencing demonstrated that SC inhibition impaired detection more strongly than V1 inhibition (Figure 4). Together, these converging results from neural recordings and causal manipulations establish that the SC is a stronger contributor to visual spatial detection than V1 in freely moving mice.

With growing interest in studying vision under naturalistic conditions (Dennis et al., 2021; Franke et al., 2024; Hoy and Farrow, 2025; Klioutchnikov et al., 2023; Qiu et al., 2021; Shapcott et al., 2025; Singh et al., 2025; Skyberg and Niell, 2024; Syeda et al., 2024), several methods have been developed to monitor eye and head movements (Meyer et al., 2020, 2018; Michaiel et al., 2020; Wallace et al., 2013), and to reconstruct visual input from world-view cameras (Parker et al., 2022). However, controlling visual input in freely moving mice has remained a significant challenge. Our closed-loop system controls visual stimuli in real time, based on the animal’s location and head angle, tracked with DeepLabCut (Kane et al., 2020; Mathis et al., 2018). This approach enables spatially and temporally controlled stimulus presentation, and evokes spatially selective visual responses in both SC and V1 (Figure 1). This capability enables the manipulation of visual features, supports repeated trials, and facilitates data analysis, allowing for validation and extension of findings from traditional head-fixed studies.

In head-fixed mice, activity in both SC and V1 has been linked to detection performance (Cazemier et al., 2024; Cone et al., 2020; Montijn et al., 2016, 2015). Using a visual spatial detection task in freely moving mice, we find that SC activity correlates more strongly with reaction time than V1 activity (Figure 2). The effect was evident at both population and single-cell levels, across fast and slow trial classifications and continuous reaction time measures (Figure 2). This supports a more critical role of SC in visual spatial detection, consistent with its established role in guiding approach and orienting behaviors (Brenner et al., 2023; Essig et al., 2021; González-Rueda et al., 2024; Hoy et al., 2019; Krauzlis et al., 2013; Masullo et al., 2019; Mysore and Knudsen, 2011; Zahler et al., 2021; Zucca et al., 2025).

A notable feature of SC firing activity was its persistence from stimulus onset until the mouse touched the stimulus, particularly for contralateral stimuli. This resembles “prelude” activity in primate SC, where neurons maintain firing until saccade initiation (Basso and May, 2017; Basso and Wurtz, 1998; Gandhi and Katnani, 2011; Glimcher and Sparks, 1992; Katz et al., 2023; McPeek and Keller, 2002; Munoz and Wurtz, 1995; Stine et al., 2023). Sustained, spatially selective SC activity has also been reported in rats during olfactory-guided tasks (Felsen and Mainen, 2008), suggesting that SC activity can generally persist until the animal receives the task outcome. In our case, SC activity terminated at the time of stimulus touch. This likely occurred because a click sound was presented at touch onset. Overall, these observations suggest that SC neurons may sustain their activity throughout ‘task engagement’ and stop firing once sensory information signals ‘goal achievement or reward expectation’ (Baruchin et al., 2023; Felsen and Mainen, 2008; Griggs et al., 2018; Stine et al., 2023). However, given differences in species, sensory modalities and task designs, further research can identify the specific contexts that drive such sustained activity.

Optogenetic silencing further established SC’s dominant role in visual spatial detection tasks. We found that SC inhibition reduced detection performance more than V1 inhibition. Both SC and V1 inhibition have previously been shown to affect detection (Cone et al., 2024; Day-Cooney et al., 2022; Glickfeld et al., 2013; Goldbach et al., 2021; Jin and Glickfeld, 2020; Poort et al., 2015; Wang et al., 2020). In our study, SC and V1 activity were recorded simultaneously in the same mouse, and inhibition conditions were interleaved within the same session. This design enabled a direct comparison of their causal contributions. The effect was spatially specific, with reduced detection for contralateral but not ipsilateral stimuli. Notably, mice could still detect stimuli under V1 inhibition, resembling blindsight in humans (Leopold, 2012). However, because mouse SC receives more direct retinal input than primate SC (Ellis et al., 2016; Hafed et al., 2023; Perry and Cowey, 1984), future studies can compare these effects in freely-moving primates.

Our findings open new directions for future investigation in freely moving mice. First, given the interactions between SC and V1 (Ahmadlou et al., 2018, 2017; Ito and Feldheim, 2018; Resulaj, 2021; Zhao et al., 2014), pathway-specific manipulations can clarify how specific projections contribute to visual spatial detection. Second, future studies can investigate the interactions between different SC layers and their role in visual spatial detection. The superficial SC (sSC) is primarily visual and the deep SC (dSC) is predominantly motor-related (Ito et al., 2017; Lee et al., 2020; Takaura et al., 2011). There are monosynaptic projections from sSC to dSC (Doubell et al., 2003) and feedback loops from motor to sensory layers (Ghitani et al., 2014).

Animals operate in a dynamic and complex world. Studying behavior in freely moving subjects enables understanding how specific neural circuits cope with the unique challenges posed by naturalistic environments. Our results reveal the distinct contributions to visual spatial detection and the distinct patterns of neural activity of two key visual processing hubs in mammals. Targeted optogenetic silencing of SC and V1 causes remarkably localized perturbations, underscoring their relative roles in visually-guided action. By integrating closed-loop visual stimulation, neural recordings, optogenetic perturbations, and eye and head tracking, we provide a framework to bridge the gap between head-fixed and freely moving studies. This approach opens up new avenues to study neural mechanisms of visually-guided decision-making and action in unrestrained animals.

## Author contributions

Conceptualization and Investigation, J.K. and J.P.; Supervision, J.P. and R.B.; Methodology, Software, and Data Curation, J.K. and J.P.; Formal Analysis and Visualization, J.K. and J.P.; Writing, Funding Acquisition, J.K., R.B., and J.P.

## Acknowledgements

We thank Edina Horvath-Gulacsi for her help with histology. We thank all members of the Poort and Beltramo labs and Arne Meyer for helpful discussions. We thank John McClure and Lilia Kukovska for help with the behavior and eye tracking.

## Funding

This work was supported by a Wolfson College & Department of Physiology, Development and Neuroscience (PDN) and School of the Biological Sciences University of Cambridge studentship (J.K.), the Wellcome Trust and the Royal Society (J.P., 211258/Z/18/Z, R.B., 222583/Z/21/Z), and a UKRI Medical Research Council Equipment Grant (MC-PC-MR-X012271/1).

## Methods

### Experimental subjects

Experiments were performed on 7 male VGAT-cre x Ai 32 mice, which were crossed from VGAT-cre (JAX:028862) and Ai 32 mice (JAX:024109). Mice were housed in reversed light/dark 12-hour cycle conditions. Mice had free access to water and food during surgery recovery. Mice were food-deprived for visual detection tasks while maintaining their weight at 85% or more of their initial baseline weight. All experimental procedures were carried out in accordance with the institutional animal welfare guidelines and with AWERB (Animal Welfare and Ethical Review Board) and UK Home Office Project License approval under the United Kingdom Animals (Scientific Procedures) Act of 1986.

### Surgical Procedures

Healthy male mice between 7-14 weeks were anesthetized with 3% isoflurane and maintained with 1-2% isoflurane with 1.5% oxygen. Metacam (5mg/kg) was injected for analgesia, and sterile saline (0.1 ml) was injected every 1-2 hours for hydration. A heating pad was used to keep body temperature at 36 - 38°C. The mouse was fixed with a stereotaxic instrument (Model 963, Kopf Instruments, USA). Eye ointment (Xailin Night) was used to protect the eyes. The head was shaved, and the scalp was cut into a circular piece to expose the skull. The headplate was attached to the skull with cement (Superbond C&B, Sun Medical, Japan). The positions of V1 (anteroposterior: −4.10 mm; mediolateral: 2.45 mm; dorsoventral: −0.3 mm) and SC (anteroposterior: −3.8 mm; mediolateral: 0.7 mm; dorsoventral: −1.2 mm) were marked with a needle. After creating two craniotomies for V1 and SC, a custom-built microdrive (Axona) was implanted. The custom-built microdrive included two optrodes, each consisting of 32-channel tetrodes for V1 and SC (64 channels total), along with two fiber optic cannula (Thorlabs, CFMLC52U-20, Fiber Optic Cannula, Ø1.25 mm Ceramic Ferrule, Ø200 µm Core, 0.50 NA). After implantation, the microdrive was protected by cement (Superbond C&B, Sun Medical, Japan). All mice were monitored for 5 days postoperatively. Experiments were started after at least 5 days of recovery.

### Head-mounted Camera System

Details of the custom head-mounted camera system have been reported in previous studies (Meyer et al., 2020, 2018). Camera modules (Adafruit Spy Camera, Adafruit, USA) were used for monitoring the eyes. A custom 3D printed camera holder (Meyer et al., 2018) with 21G cannula (Coopers Needle Works, UK) was used to hold the camera, IR LEDs (VSMB2943GX01, Vishay, USA), and a 7.0 mm x 9.3 mm IR mirror (Calflex-X NIR-Blocking Filter, Optics Balzers, Germany). A connector (852-10-00810-001101, Preci-Dip, Switzerland) was used to attach the camera holder to the headplate of the mice. The mice were head-fixed, and the mirror position was modified until the eye was in the center of the eye cameras. Epoxy (Araldite Rapid, Araldite, UK) was used to fix the mirror position. Single-board computers (Raspberry Pi 3 model B, Raspberry Pi Foundation, UK) were used for recording camera data. The cameras were collected at 60 Hz. An IMU sensor (MPU-9250, InvenSense, USA) was used to measure head roll, pitch, and yaw. A custom Open-Ephys plugin was used to record the sensor data.

### Visual Task Experiment Chamber Set-up

To study visual detection in freely behaving mice, we used a 12.1 inch touchscreen (NEX121, Nexio, Korea). The training chamber had a trapezoidal shape (width of box at screen side: 24 cm, opposite side 6 cm). Custom Python software on a single-board computer (Raspberry Pi 3B, Raspberry Pi Foundation, UK) was used to present the visual stimuli and detect touchscreen presses. A lick spout was located on the opposite side of the touchscreen, inside an IR beam break detector detecting nose pokes when the animal licked (OPB815WZ, TT Electronics, UK). A pinch valve (161P011, NResearch, USA) controlled by a valve driver (CDS-V01, NResearch, USA) was connected to silicon tubing (TBGM100, NResearch, USA) attached to the lick spout to give strawberry milk rewards.

### Pupil Position Tracking Using DLC

Manually selected 40-60 frames from the training video were labelled with 13 detailed parts, including eye position (eye top, eye bottom, eye center, eye left, eye right), and pupil position (pupil position 1 to 8). We trained a deep convolutional network using DeepLabCut (Mathis et al., 2018). The ResNet101 model was used and trained with 150,000-500,000 iterations. Eye videos were analyzed with the trained network. The predicted positions of eye parts were extracted and used for pupil position analysis.

### Mouse Position Tracking Using DLC

Styrofoam markers were covered with reflective tape. Two different patterns (dots and stripes) were painted on the reflective tape with black paint (Black 3.0, Stuart Semple, UK). Sample videos were recorded after attaching the styrofoam markers to mice. Manually selected 40-60 frames were labelled with 9 detailed parts, including training box positions (screen left, screen right, reward left, reward right), mouse marker positions (left marker, right marker), and mouse body components (mouse head, mouse center, mouse tail). We used a ResNet101 network for training using DeepLabCut (Mathis et al., 2018). After training, the model was used for real-time tracking using DeepLabCut-live to trigger stimulus presentation (Kane et al., 2020).

### Optogenetic inhibition of V1 and SC

We used VGAT-Cre x Ai32 mice, where Channelrhodopsin-2 (ChR2) is expressed in VGAT-expressing cells, enabling us to stimulate ChR2-expressing GABAergic neurons using a blue LED (Thorlabs, M470F3 - 470 nm fiber-coupled LED). Two LEDs were connected to the fiber optic cannula (Thorlabs, CFMLC52U-20, Ø1.25 mm Ceramic Ferrule, Ø200 µm Core, 0.50 NA) in each area (V1, SC), with their maximum output power measured at the fiber tip of 3 mW.

### Visual Detection Task

The mice were first habituated to the experimental chamber for 10 minutes each day over 1–2 days. Following habituation, they were trained to drink strawberry milk from the lick spout. A large drifting grating stimulus (25 x 25 cm, contrast: 100%) was presented, and the mice were rewarded if they touched the stimulus within 30 seconds of its onset. Once they successfully completed 30 consecutive trials, the target size was progressively reduced to 15 x 15 cm and then to 7 x 7 cm. During this training period, the grating stimulus was presented only in the center of the screen. After the mice successfully performed the detection task at one location for 30 consecutive trials, a drifting grating stimulus was presented at three random locations (left: −7.0 cm; center: 0 cm; right: +7.0 cm). Once the mice successfully completed the detection task across these three random locations, visual detection tasks with optogenetic inhibition were conducted. For the visual detection experiments with optogenetic inhibition, LEDs were turned on at stimulus onset for 4 seconds. A total of 9 conditions were analyzed, defined by three stimulus locations (left, center, and right) and three optogenetic conditions: control without optogenetic manipulation, V1 inhibition, and SC inhibition. These conditions were randomly interleaved across trials.

### Electrophysiological Recordings

Two 32-channel Intan RHD 2132 amplifier boards (Intan Technologies, USA) and two flexible serial peripheral interface cables (“Ultra Thin RHD2000 SPI Cable,” Intan Technologies, USA) were used for in vivo extracellular recording in V1 and SC. An Open Ephys acquisition board (Open Ephys) was used to acquire data from 64 channels, which were digitized at 30 kHz. Once the mice had recovered from microdrive implantation surgery, each tetrode in V1 and SC was gradually lowered by 10–15 µm per day until cell spikes were detected. After the visual detection experiment, each tetrode was further lowered by 20–50 µm until additional cell spikes were detected. Recordings were obtained from each mouse at 3–10 different depths within the following ranges: V1, dorsoventral −300 to −700 µm; sSC, dorsoventral −1200 to −1500 µm. MATLAB was used for offline data analysis. Electrophysiological recordings from V1 and SC were processed using Kilosort 2.0 (Pachitariu et al., 2024) to detect spikes. Single units were defined as those with firing rates between 0.5 Hz and 50 Hz and an inter-spike interval (2 ms) contamination rate below 1%. We used PHY 2.0 (https://github.com/cortex-lab/phy) to review all units and exclude outliers (noise or spikes from other units) using a Mahalanobis distance threshold of 10 (Hill et al., 2011).

### Histology

Histology was performed similarly to previous studies (Gadiwalla et al., 2025). Mice were anaesthetized by intraperitoneal injection of pentobarbital and perfused transcardially with PBS (15 ml) with heparin (102 mg/L) and 10 ml paraformaldehyde (PFA, 4%) in PIPES buffer (60 mM PIPES, 25 mM HEPES, 5 mM EGTA, and 1 mM MgCl2). Brains were extracted and postfixed in 4% PFA for 24 hours and then placed into PBS/Azide (0.02% NaN3). Brains were embedded in 5% agarose and coronal sections sliced around V1 and SC with 100µm thickness using a vibratome (VT1000S, Leica). Slices were then washed in PBS and mounted on glass slides (Menzel-Gläser) with mounting medium with DAPI stain (AB104139, Abcam). The electrode location was confirmed using bright-field images (see Figure S1G-H), as in previous studies (Beltramo and Scanziani, 2019).

## Declaration of interests and use of AI

The authors declare no competing interests. During the preparation of this work, J.K. used Grammarly and ChatGPT to check grammar and spelling. After using this tool, the authors reviewed and edited the content as needed and take responsibility for the content of the publication.

## Supplementary information

**Figure S1.**
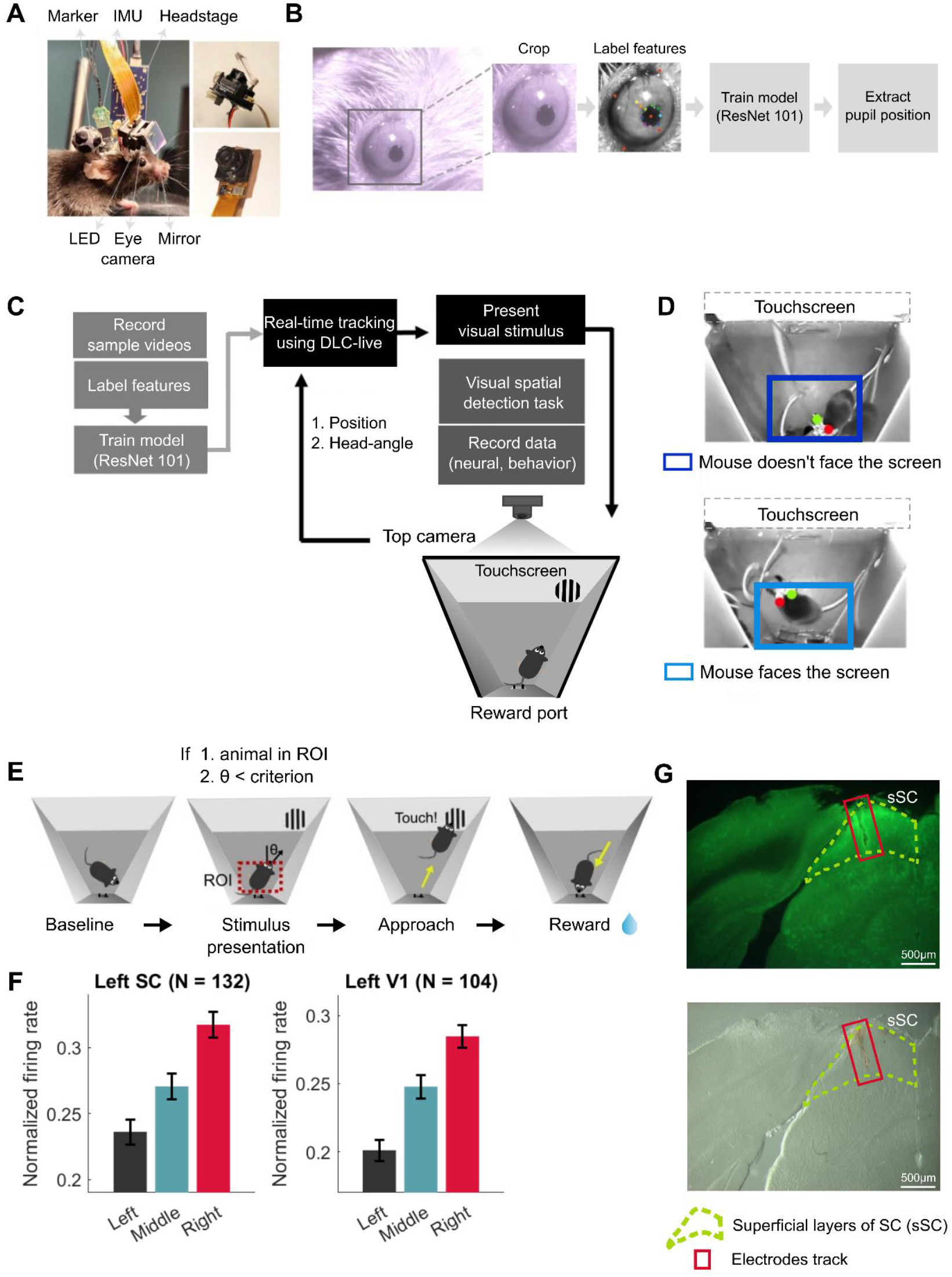
Closed-loop visual stimulus presentation and an integrated system to monitor behavior and neural activity in SC and V1. Related to Figure 1. **(A)** Integrated neural recording system consisting of an infrared LED, an infrared mirror, an eye camera, an inertial motion sensor (IMU), styrofoam markers, and drive mechanisms for neural recording from SC and V1. **(B)** Pupil position tracking using DeepLabCut (DLC). A region of interest (ROI) was selected. A total of 13 features (eye and pupil points) were manually labeled, and a ResNet-101 model was trained to extract feature positions. **(C)** Closed-loop visual stimulus presentation system using DLC-live. Videos were trained using ResNet-101, and the pre-trained model was applied to detect mouse position and head angle in real time. **(D)** Real-time mouse tracking using DLC-live. Dots (green and red) represent styrofoam markers (green: dot pattern; red: stripe pattern). Squares indicate ROIs. Top: the mouse is in the ROI but not facing the screen. Bottom: the mouse is within the ROI and facing the screen. Stimuli were presented only when the mouse was in the ROI and its absolute head angle relative to the touchscreen was less than 45° (0° indicates that the mouse was directly facing the touchscreen). **(E)** Schematic of the visual detection task. When the mouse was within the ROI and its head angle was less than 45°, the closed-loop system presented a visual stimulus. The mouse then approached and touched the stimulus. If the mouse successfully touched the stimulus within 30 seconds, a strawberry milk reward was delivered at the lick spout. **(F)** Normalized firing rate of SC and V1 in response to the three stimulus locations. Stimulus-evoked firing activity was measured from 0 to 1 second. Error bars, SEM. **(G)** Histology showing electrode tracks in SC and V1 of a VGAT-Cre × Ai32 mouse. The green dotted line indicates the superficial SC layer. The red box marks the electrode tracks. Top: green fluorescence image. Bottom: brightfield contrast image.

**Figure S2.**
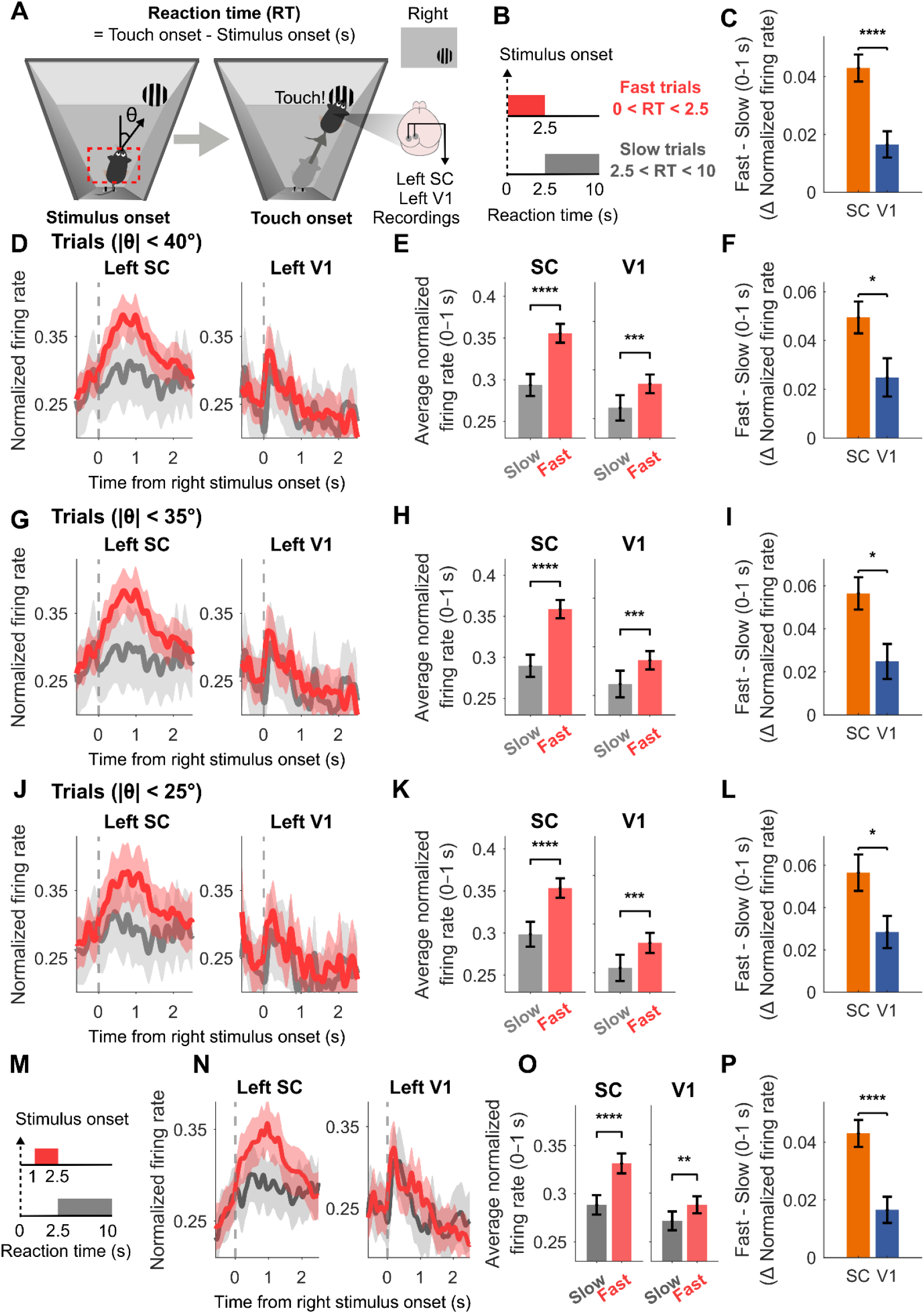
SC activity shows greater difference between reaction time conditions than V1 across head-angle and behavioral control analyses. Related to Figure 2. **(A)** Schematic of the visual spatial detection task. Reaction time (RT) was defined as the duration from the stimulus onset to the touch. A right stimulus was presented at stimulus onset, and the touch was the time when the mouse first touched the right stimulus. **(B)** Trials were classified into two classes: Fast trials (0 < RT < 2.5 s) and Slow trials (2.5 < RT < 10 s). (C) The difference between Fast and Slow trials in SC and V1. Mean firing rates from 0-1 s were calculated for both classes, and the difference was computed as Fast - Slow (Wilcoxon rank-sum test, p = 0.0019). Error bars, SEM. **(D-L)** Control analyses testing the effect of head angle on SC and V1 firing activity. Only trials with head angles below specific criteria (40°, 35°, or 25°) were included in each analysis. **(D-F)** Results with head-angle restricted to <40°. (D) Normalized firing rates of SC and V1. (Slow: gray, Fast: red). Shaded area represents SEM. (E) Average neural activity (0–1 s). Error bars, SEM. (F) The difference between Fast and Slow trials in SC and V1. Error bars, SEM. SC showed a significantly greater difference between classes (Wilcoxon rank-sum test, p = 0.013). **(G-I)** Results with head-angle restricted to <35°. (G) Normalized firing rates of SC and V1. (H) Average neural activity (0–1 s). (I) SC showed a significantly greater difference between classes (Wilcoxon rank-sum test, p = 0.011). **(J-L)** Results with head-angle restricted to <25°. (J) Normalized firing rates of SC and V1. (K) Average neural activity (0–1 s). (L) SC showed a significantly greater difference between classes (Wilcoxon rank-sum test, p = 0.018). **(M-P)** SC and V1 neural responses across behavioral classes were defined as Fast (1 < RT < 2.5 s) and Slow (2.5 < RT < 10 s), excluding trials with RT < 1 s. (N) Normalized firing rates of SC and V1 across classes. (O) Average neural activity (0–1 s). (P) The difference between Fast and Slow trials. Error bars, SEM. SC showed a significantly greater difference than V1 (Wilcoxon rank-sum test, p = 8.1 × 10^−5^).

**Figure S3.**
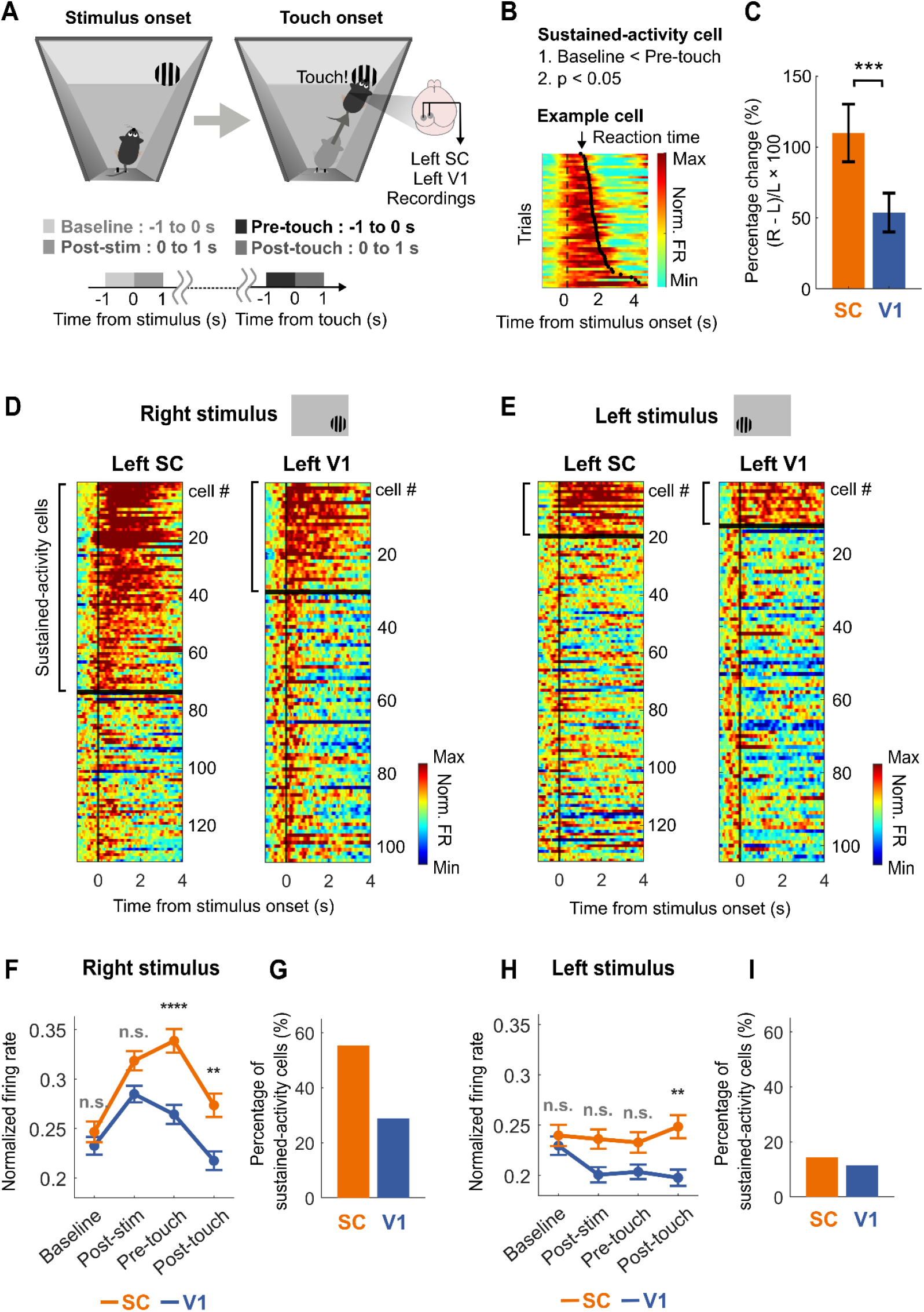
SC showed more spatially selective sustained activity compared to V1. **(A)** Schematic of the visual spatial detection task. Four phases were defined throughout the detection task: Baseline (−1 to 0 s before stimulus onset), Post-stimulus (0 to 1 s after stimulus onset), Pre-touch (−1 to 0 s before touch onset), and Post-touch (0 to 1 s after touch onset). **(B)** Sustained cells were identified as those with significantly higher Pre-touch firing activity relative to Baseline firing activity. (C) Percentage difference between left and right stimulus firing rates during the Pre-touch period, calculated as (Right - Left) / Left × 100. Error bars, SEM. **(D-E)** SC and V1 firing activity in response to the right (D) and left (E) stimuli. The mean baseline firing activity (−1 to 0 s) was subtracted from normalized firing rates for each cell. Cells were sorted by their Pre-touch/Baseline ratio. Horizontal black lines mark the boundary between sustained and non-sustained cells. Sustained cells were defined as showing significantly higher Pre-touch firing activity than Baseline firing activity. **(F)**Temporal dynamics of SC and V1 in response to the right stimulus. **(G)** Percentage of sustained cells in SC and V1 in response to the right stimulus (SC: 55.3%, V1: 28.4%). **(H)**Temporal dynamics of SC and V1 in response to the left stimulus. **(I)**Percentage of sustained cells in SC and V1 in response to the left stimulus (SC: 14.4%, V1: 11.5%).

**Figure S4.**
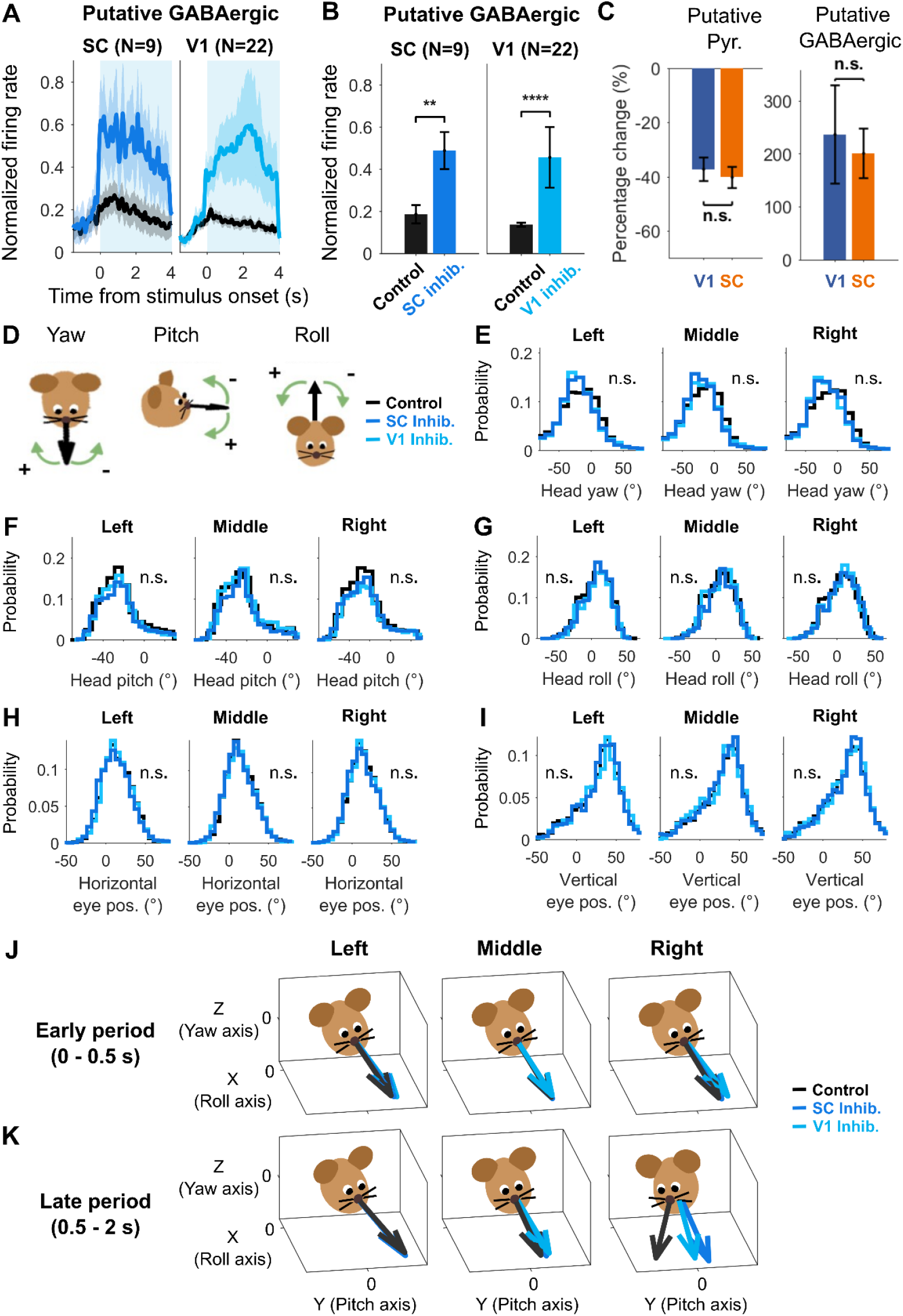
Impact of optogenetic inhibition on neural activity, head orientation, and eye position. Related to Figure 4. (A) Putative GABAergic cell activity during local optogenetic stimulation in V1 (left) and the SC (right). In each panel, the black trace shows activity under the control condition, and the blue trace shows activity under the corresponding local inhibition condition (SC inhibition for the left panel, V1 inhibition for the right panel). Shaded area represents SEM. (B) Mean firing activity (0-4 s) of putative GABAergic cells during local optogenetic stimulation in V1 (left) and the SC (right). Error bars, SEM. Local optogenetic stimulation significantly increased putative GABAergic cell activity (SC: p = 3.91 × 10^− 3^, V1: p = 3.09 × 10^−5^; Wilcoxon signed rank test). (C) Percentage change from control to inhibition for putative pyramidal cells (left) and putative GABAergic cells (right), computed as (Control − Inhibition) / Control × 100. Error bars, SEM. No significant difference was found between V1 and SC (putative pyramidal: p = 0.48; putative GABAergic: p = 0.53; Wilcoxon rank-sum test). (D) Schematic illustrating head yaw, pitch, and roll. (E–I) Head orientation (yaw, pitch, roll) and eye position (x, y) during the early phase (0–0.5 s) across conditions: control (black), V1 inhibition (light blue), and SC inhibition (blue). No significant differences were observed between control and V1 inhibition or between control and SC inhibition (Wilcoxon signed-rank test, Bonferroni-corrected; p > 0.05). (J) Head orientation during visual detection in the early period. No significant differences were observed between control and V1 inhibition or between control and SC inhibition (Wilcoxon signed-rank test, Bonferroni-corrected; p > 0.05). (K) Head orientation during visual detection in the late period (0.5–2 s). Significant head-yaw changes were observed only for right-side stimuli (left: p > 0.05; middle: p > 0.05; right: control vs V1 inhibition, p = 4.28 × 10^−7^; control vs SC inhibition, p = 3.17 × 10^−7^; V1 inhibition vs SC inhibition, p = 0.0093; Wilcoxon signed-rank test, Bonferroni-corrected).

## Notes

### Competing Interest Statement

The authors have declared no competing interest.

## References

Ahmadlou, M., Tafreshiha, A., Heimel, J.A., 2017. Visual Cortex Limits Pop-Out in the Superior Colliculus of Awake Mice. Cereb. Cortex 27, 5772–5783. 10.1093/cercor/bhx254

Ahmadlou, M., Zweifel, L.S., Heimel, J.A., 2018. Functional modulation of primary visual cortex by the superior colliculus in the mouse. Nat. Commun. 9, 3895. 10.1038/s41467-018-06389-6

Baruchin, L.J., Alleman, M., Schröder, S., 2023. Reward Modulates Visual Responses in the Superficial Superior Colliculus of Mice. J. Neurosci. 43, 8663–8680. 10.1523/JNEUROSCI.0089-23.2023

Basso, M.A., Bickford, M.E., Cang, J., 2021. Unraveling circuits of visual perception and cognition through the superior colliculus. Neuron 109, 918–937. 10.1016/j.neuron.2021.01.013

Basso, M.A., May, P.J., 2017. Circuits for Action and Cognition: A View from the Superior Colliculus. Annu. Rev. Vis. Sci. 3, 197–226. 10.1146/annurev-vision-102016-061234

Basso, M.A., Wurtz, R.H., 1998. Modulation of Neuronal Activity in Superior Colliculus by Changes in Target Probability. J. Neurosci. 18, 7519–7534. 10.1523/JNEUROSCI.18-18-07519.1998

Beltramo, R., 2020. A new primary visual cortex. Science 370, 46–46. 10.1126/science.abe1482

Beltramo, R., Scanziani, M., 2019. A collicular visual cortex: Neocortical space for an ancient midbrain visual structure. Science 363, 64–69. 10.1126/science.aau7052

Brenner, J.M., Beltramo, R., Gerfen, C.R., Ruediger, S., Scanziani, M., 2023. A genetically defined tectothalamic pathway drives a system of superior-colliculus-dependent visual cortices. Neuron S0896627323003070. 10.1016/j.neuron.2023.04.022

Cazemier, J.L., Haak, R., Tran, T.L., Hsu, A.T., Husic, M., Peri, B.D., Kirchberger, L., Self, M.W., Roelfsema, P., Heimel, J.A., 2024. Involvement of superior colliculus in complex figure detection of mice. eLife 13, e83708. 10.7554/eLife.83708

Cone, J.J., Bade, M.L., Masse, N.Y., Page, E.A., Freedman, D.J., Maunsell, J.H.R., 2020. Mice Preferentially Use Increases in Cerebral Cortex Spiking to Detect Changes in Visual Stimuli. J. Neurosci. 40, 7902–7920. 10.1523/JNEUROSCI.1124-20.2020

Cone, J.J., Mitchell, A.O., Parker, R.K., Maunsell, J.H.R., 2024. Stimulus-dependent differences in cortical versus subcortical contributions to visual detection in mice. Curr. Biol. 34, 1940-1952.e5. 10.1016/j.cub.2024.03.061

Day-Cooney, J., Cone, J.J., Maunsell, J.H.R., 2022. Perceptual Weighting of V1 Spikes Revealed by Optogenetic White Noise Stimulation. J. Neurosci. 42, 3122–3132. 10.1523/JNEUROSCI.1736-21.2022

Dennis, E.J., El Hady, A., Michaiel, A., Clemens, A., Tervo, D.R.G., Voigts, J., Datta, S.R., 2021. Systems Neuroscience of Natural Behaviors in Rodents. J. Neurosci. 41, 911–919. 10.1523/JNEUROSCI.1877-20.2020

Doubell, T.P., Skaliora, I., Baron, J., King, A.J., 2003. Functional Connectivity between the Superficial and Deeper Layers of the Superior Colliculus: An Anatomical Substrate for Sensorimotor Integration. J. Neurosci. 23, 6596–6607. 10.1523/JNEUROSCI.23-16-06596.2003

Ellis, E.M., Gauvain, G., Sivyer, B., Murphy, G.J., 2016. Shared and distinct retinal input to the mouse superior colliculus and dorsal lateral geniculate nucleus. J. Neurophysiol. 116, 602–610. 10.1152/jn.00227.2016

Essig, J., Hunt, J.B., Felsen, G., 2021. Inhibitory neurons in the superior colliculus mediate selection of spatially-directed movements. Commun. Biol. 4, 719. 10.1038/s42003-021-02248-1

Felsen, G., Mainen, Z.F., 2008. Neural Substrates of Sensory-Guided Locomotor Decisions in the Rat Superior Colliculus. Neuron 60, 137–148. 10.1016/j.neuron.2008.09.019

Franke, K., Cai, C., Ponder, K., Fu, J., Sokoloski, S., Berens, P., Tolias, A.S., 2024. Asymmetric distribution of color-opponent response types across mouse visual cortex supports superior color vision in the sky. eLife 12, RP89996. 10.7554/eLife.89996

Gadiwalla, S., Guillaume, C., Huang, L., White, S.J.B., Basha, N., Petersen, P.H., Galliano, E., 2025. Ex Vivo Functional Characterization of Mouse Olfactory Bulb Projection Neurons Reveals a Heterogeneous Continuum. eneuro 12, ENEURO.0407-24.2025. 10.1523/ENEURO.0407-24.2025

Gandhi, N.J., Katnani, H.A., 2011. Motor Functions of the Superior Colliculus. Annu. Rev. Neurosci. 34, 205–231. 10.1146/annurev-neuro-061010-113728

Gantar, L., Burgess, M.A., Mansour, N., Rusco-Portabella, J., Lowe, D.M.T., Námešná, A., Gill, D., Harris, I., Orlowska-Feuer, P., Ebrahimi, A.S., Storchi, R., Petersen, R.S., 2025. Encoding of body state in whisker-related somatosensory cortex of freely moving mice. Curr. Biol. 35, 3461-3472.e5. 10.1016/j.cub.2025.06.040

Ghitani, N., Bayguinov, P.O., Vokoun, C.R., McMahon, S., Jackson, M.B., Basso, M.A., 2014. Excitatory Synaptic Feedback from the Motor Layer to the Sensory Layers of the Superior Colliculus. J. Neurosci. 34, 6822–6833. 10.1523/JNEUROSCI.3137-13.2014

Glickfeld, L.L., Histed, M.H., Maunsell, J.H.R., 2013. Mouse Primary Visual Cortex Is Used to Detect Both Orientation and Contrast Changes. J. Neurosci. 33, 19416–19422. 10.1523/JNEUROSCI.3560-13.2013

Glimcher, P.W., Sparks, D.L., 1992. Movement selection in advance of action in the superior colliculus. Nature 355, 542–545. 10.1038/355542a0

Goldbach, H.C., Akitake, B., Leedy, C.E., Histed, M.H., 2021. Performance in even a simple perceptual task depends on mouse secondary visual areas. eLife 10, e62156. 10.7554/eLife.62156

González-Rueda, A., Jensen, K., Noormandipour, M., De Malmazet, D., Wilson, J., Ciabatti, E., Kim, J., Williams, E., Poort, J., Hennequin, G., Tripodi, M., 2024. Kinetic features dictate sensorimotor alignment in the superior colliculus. Nature 631, 378–385. 10.1038/s41586-024-07619-2

Griggs, W.S., Amita, H., Gopal, A., Hikosaka, O., 2018. Visual Neurons in the Superior Colliculus Discriminate Many Objects by Their Historical Values. Front. Neurosci. 12, 396. 10.3389/fnins.2018.00396

Hafed, Z.M., Hoffmann, K.-P., Chen, C.-Y., Bogadhi, A.R., 2023. Visual Functions of the Primate Superior Colliculus. Annu. Rev. Vis. Sci. 9, 361–383. 10.1146/annurev-vision-111022-123817

Hill, D.N., Mehta, S.B., Kleinfeld, D., 2011. Quality Metrics to Accompany Spike Sorting of Extracellular Signals. J. Neurosci. 31, 8699–8705. 10.1523/JNEUROSCI.0971-11.2011

Hoy, J.L., Bishop, H.I., Niell, C.M., 2019. Defined Cell Types in Superior Colliculus Make Distinct Contributions to Prey Capture Behavior in the Mouse. Curr. Biol. 29, 4130-4138.e5. 10.1016/j.cub.2019.10.017

Hoy, J.L., Farrow, K., 2025. The superior colliculus. Curr. Biol. 35, R164–R168. 10.1016/j.cub.2025.01.022

Ito, S., Feldheim, D.A., 2018. The Mouse Superior Colliculus: An Emerging Model for Studying Circuit Formation and Function. Front. Neural Circuits 12, 10. 10.3389/fncir.2018.00010

Ito, S., Feldheim, D.A., Litke, A.M., 2017. Segregation of Visual Response Properties in the Mouse Superior Colliculus and Their Modulation during Locomotion. J. Neurosci. 37, 8428–8443. 10.1523/JNEUROSCI.3689-16.2017

Jin, M., Glickfeld, L.L., 2020. Mouse Higher Visual Areas Provide Both Distributed and Specialized Contributions to Visually Guided Behaviors. Curr. Biol. 30, 4682-4692.e7. 10.1016/j.cub.2020.09.015

Kane, G.A., Lopes, G., Saunders, J.L., Mathis, A., Mathis, M.W., 2020. Real-time, low-latency closed-loop feedback using markerless posture tracking. eLife 9, e61909. 10.7554/eLife.61909

Katz, L.N., Yu, G., Herman, J.P., Krauzlis, R.J., 2023. Correlated variability in primate superior colliculus depends on functional class. Commun. Biol. 6, 540. 10.1038/s42003-023-04912-0

Klioutchnikov, A., Wallace, D.J., Sawinski, J., Voit, K.-M., Groemping, Y., Kerr, J.N.D., 2023. A three-photon head-mounted microscope for imaging all layers of visual cortex in freely moving mice. Nat. Methods 20, 610–616. 10.1038/s41592-022-01688-9

Krauzlis, R.J., Lovejoy, L.P., Zénon, A., 2013. Superior Colliculus and Visual Spatial Attention. Annu. Rev. Neurosci. 36, 165–182. 10.1146/annurev-neuro-062012-170249

Lanzarini, F., Maranesi, M., Rondoni, E.H., Albertini, D., Ferretti, E., Lanzilotto, M., Micera, S., Mazzoni, A., Bonini, L., 2025. Neuroethology of natural actions in freely moving monkeys. Science 387, 214–220. 10.1126/science.adq6510

Lee, K.H., Tran, A., Turan, Z., Meister, M., 2020. The sifting of visual information in the superior colliculus. eLife 9, e50678. 10.7554/eLife.50678

Leopold, D.A., 2012. Primary Visual Cortex: Awareness and Blindsight. Annu. Rev. Neurosci. 35, 91– 109. 10.1146/annurev-neuro-062111-150356

Liu, B., Huberman, A.D., Scanziani, M., 2016. Cortico-fugal output from visual cortex promotes plasticity of innate motor behaviour. Nature 538, 383–387. 10.1038/nature19818

Masullo, L., Mariotti, L., Alexandre, N., Freire-Pritchett, P., Boulanger, J., Tripodi, M., 2019. Genetically Defined Functional Modules for Spatial Orienting in the Mouse Superior Colliculus. Curr. Biol. 29, 2892-2904.e8. 10.1016/j.cub.2019.07.083

Mathis, A., Mamidanna, P., Cury, K.M., Abe, T., Murthy, V.N., Mathis, M.W., Bethge, M., 2018. DeepLabCut: markerless pose estimation of user-defined body parts with deep learning. Nat. Neurosci. 21, 1281–1289. 10.1038/s41593-018-0209-y

McPeek, R.M., Keller, E.L., 2002. Saccade Target Selection in the Superior Colliculus During a Visual Search Task. J. Neurophysiol. 88, 2019–2034. 10.1152/jn.2002.88.4.2019

Meyer, A.F., O’Keefe, J., Poort, J., 2020. Two Distinct Types of Eye-Head Coupling in Freely Moving Mice. Curr. Biol. 30, 2116-2130.e6. 10.1016/j.cub.2020.04.042

Meyer, A.F., Poort, J., O’Keefe, J., Sahani, M., Linden, J.F., 2018. A Head-Mounted Camera System Integrates Detailed Behavioral Monitoring with Multichannel Electrophysiology in Freely Moving Mice. Neuron 100, 46-60.e7. 10.1016/j.neuron.2018.09.020

Michaiel, A.M., Abe, E.T., Niell, C.M., 2020. Dynamics of gaze control during prey capture in freely moving mice. eLife 9, e57458. 10.7554/eLife.57458

Montijn, J.S., Goltstein, P.M., Pennartz, C.M., 2015. Mouse V1 population correlates of visual detection rely on heterogeneity within neuronal response patterns. eLife 4, e10163. 10.7554/eLife.10163

Montijn, J.S., Olcese, U., Pennartz, C.M.A., 2016. Visual Stimulus Detection Correlates with the Consistency of Temporal Sequences within Stereotyped Events of V1 Neuronal Population Activity. J. Neurosci. 36, 8624–8640. 10.1523/JNEUROSCI.0853-16.2016

Munoz, D.P., Wurtz, R.H., 1995. Saccade-related activity in monkey superior colliculus. I. Characteristics of burst and buildup cells. J. Neurophysiol. 73, 2313–2333. 10.1152/jn.1995.73.6.2313

Mysore, S.P., Knudsen, E.I., 2011. The role of a midbrain network in competitive stimulus selection. Curr. Opin. Neurobiol. 21, 653–660. 10.1016/j.conb.2011.05.024

Pachitariu, M., Sridhar, S., Pennington, J., Stringer, C., 2024. Spike sorting with Kilosort4. Nat. Methods 21, 914–921. 10.1038/s41592-024-02232-7

Parker, P.R.L., Abe, E.T.T., Leonard, E.S.P., Martins, D.M., Niell, C.M., 2022. Joint coding of visual input and eye/head position in V1 of freely moving mice. Neuron 110, 3897-3906.e5. 10.1016/j.neuron.2022.08.029

Parker, P.R.L., Martins, D.M., Leonard, E.S.P., Casey, N.M., Sharp, S.L., Abe, E.T.T., Smear, M.C., Yates, J.L., Mitchell, J.F., Niell, C.M., 2023. A dynamic sequence of visual processing initiated by gaze shifts. Nat. Neurosci. 26, 2192–2202. 10.1038/s41593-023-01481-7

Payne, H.L., Raymond, J.L., 2017. Magnetic eye tracking in mice. eLife 6, e29222. 10.7554/eLife.29222

Perry, V.H., Cowey, A., 1984. Retinal ganglion cells that project to the superior colliculus and pretectum in the macaque monkey. Neuroscience 12, 1125–1137. 10.1016/0306-4522(84)90007-1

Poort, J., Khan, A.G., Pachitariu, M., Nemri, A., Orsolic, I., Krupic, J., Bauza, M., Sahani, M., Keller, G.B., Mrsic-Flogel, T.D., Hofer, S.B., 2015. Learning Enhances Sensory and Multiple Non-sensory Representations in Primary Visual Cortex. Neuron 86, 1478–1490. 10.1016/j.neuron.2015.05.037

Qiu, Y., Zhao, Z., Klindt, D., Kautzky, M., Szatko, K.P., Schaeffel, F., Rifai, K., Franke, K., Busse, L., Euler, T., 2021. Natural environment statistics in the upper and lower visual field are reflected in mouse retinal specializations. Curr. Biol. 31, 3233-3247.e6. 10.1016/j.cub.2021.05.017

Resulaj, A., 2021. Projections of the Mouse Primary Visual Cortex. Front. Neural Circuits 15, 751331. 10.3389/fncir.2021.751331

Resulaj, A., Ruediger, S., Olsen, S.R., Scanziani, M., 2018. First spikes in visual cortex enable perceptual discrimination. eLife 7, e34044. 10.7554/eLife.34044

Sawinski, J., Wallace, D.J., Greenberg, D.S., Grossmann, S., Denk, W., Kerr, J.N.D., 2009. Visually evoked activity in cortical cells imaged in freely moving animals. Proc. Natl. Acad. Sci. 106, 19557–19562. 10.1073/pnas.0903680106

Senzai, Y., Fernandez-Ruiz, A., Buzsáki, G., 2019. Layer-Specific Physiological Features and Interlaminar Interactions in the Primary Visual Cortex of the Mouse. Neuron 101, 500-513.e5. 10.1016/j.neuron.2018.12.009

Shapcott, K.A., Weigand, M., Glukhova, M., Havenith, M.N., Schölvinck, M.L., 2025. DomeVR: Immersive virtual reality for primates and rodents. PLOS ONE 20, e0308848. 10.1371/journal.pone.0308848

Sharp, L., Martins, D.M., Jones, K., Niell, C.M., Neural dynamics in superior colliculus of freely moving mice. bioRxiv. 10.1101/2025.04.16.648828

Singh, V.P., Li, J., Dawson, K., Mitchell, J.F., Miller, C.T., 2025. Active vision in freely moving marmosets using head-mounted eye tracking. Proc. Natl. Acad. Sci. 122, e2412954122. 10.1073/pnas.2412954122

Skyberg, R.J., Niell, C.M., 2024. Natural visual behavior and active sensing in the mouse. Curr. Opin. Neurobiol. 86, 102882. 10.1016/j.conb.2024.102882

Stine, G.M., Trautmann, E.M., Jeurissen, D., Shadlen, M.N., 2023. A neural mechanism for terminating decisions. Neuron 111, 2601-2613.e5. 10.1016/j.neuron.2023.05.028

Syeda, A., Zhong, L., Tung, R., Long, W., Pachitariu, M., Stringer, C., 2024. Facemap: a framework for modeling neural activity based on orofacial tracking. Nat. Neurosci. 27, 187–195. 10.1038/s41593-023-01490-6

Takaura, K., Yoshida, M., Isa, T., 2011. Neural Substrate of Spatial Memory in the Superior Colliculus after Damage to the Primary Visual Cortex. J. Neurosci. 31, 4233–4241. 10.1523/JNEUROSCI.5143-10.2011

Wallace, D.J., Greenberg, D.S., Sawinski, J., Rulla, S., Notaro, G., Kerr, J.N.D., 2013. Rats maintain an overhead binocular field at the expense of constant fusion. Nature 498, 65–69. 10.1038/nature12153

Wallace, D.J., Kerr, J.N.D., 2019. Circuit interrogation in freely moving animals. Nat. Methods 16, 9–11. 10.1038/s41592-018-0275-9

Wang, L., McAlonan, K., Goldstein, S., Gerfen, C.R., Krauzlis, R.J., 2020. A Causal Role for Mouse Superior Colliculus in Visual Perceptual Decision-Making. J. Neurosci. 40, 3768–3782. 10.1523/JNEUROSCI.2642-19.2020

Wang, L., Sarnaik, R., Rangarajan, K., Liu, X., Cang, J., 2010. Visual Receptive Field Properties of Neurons in the Superficial Superior Colliculus of the Mouse. J. Neurosci. 30, 16573–16584. 10.1523/JNEUROSCI.3305-10.2010

Wheatcroft, T., Saleem, A.B., Solomon, S.G., 2022. Functional Organisation of the Mouse Superior Colliculus. Front. Neural Circuits 16, 792959. 10.3389/fncir.2022.792959

White, B.J., Kan, J.Y., Levy, R., Itti, L., Munoz, D.P., 2017. Superior colliculus encodes visual saliency before the primary visual cortex. Proc. Natl. Acad. Sci. 114, 9451–9456. 10.1073/pnas.1701003114

Zahler, S.H., Taylor, D.E., Wong, J.Y., Adams, J.M., Feinberg, E.H., 2021. Superior colliculus drives stimulus-evoked directionally biased saccades and attempted head movements in head-fixed mice. eLife 10, e73081. 10.7554/eLife.73081

Zhao, X., Liu, M., Cang, J., 2014. Visual Cortex Modulates the Magnitude but Not the Selectivity of Looming-Evoked Responses in the Superior Colliculus of Awake Mice. Neuron 84, 202–213. 10.1016/j.neuron.2014.08.037

Zucca, S., Schulz, A., Gonçalves, P.J., Macke, J.H., Saleem, A.B., Solomon, S.G., 2025. Visual loom caused by self-movement or object-movement elicits distinct responses in mouse superior colliculus. Curr. Biol. S0960982225008796. 10.1016/j.cub.2025.07.013

